# Alcohol Use Disorder Affects Brain Metals

**DOI:** 10.64898/2026.01.06.697957

**Authors:** Byron C. Jones, Wenyuan Zhao, Julia Stevens, Greg T Sutherland

**Affiliations:** Department of Genetics, Genomics, and Informatics, College of Medicine, University of Tennessee Health Science Center, Memphis, Tennessee; Department of Pharmacology, Addiction Science, and Toxicology, College of Medicine, University of Tennessee Health Science Center, Memphis, Tennessee, USA; New South Wales Brain Tissue Resource Centre, Charles Perkins Centre and School of Medical Sciences, Faculty of Medicine and Health, The University of Sydney, Sydney, NSW, Australia

**Keywords:** substantia nigra, caudate nucleus, iron, copper, zinc

## Abstract

**Background:** One of the risks associated with alcohol use disorder (AUD) is dementia. While the exact pathology is unknown, there is increasing evidence that alcohol consumption may dysregulate iron homeostasis in the brain leading to increased brain iron concentrations. Unregulated iron can produce oxidative stress and cellular damage to neurons. Iron-related cellular death is a relatively new finding and is called ferroptosis. To test the hypothesis that alcohol use disorder may cause brain iron dysregulation, we measured iron in hippocampus, caudate and substantia nigra in brain tissues obtained from individuals diagnosed with (AUD) and from individuals with no apparent alcohol-related problems.

**Methods:** We obtained hippocampus, caudate, and substantia nigra tissues from the New South Wales Brain Tissue Resource Centre, University of Sydney. The tissues came from 20 males and 20 females, half of each diagnosed with AUD. The tissues were wet-ashed and prepared for analysis of iron, copper, and zinc by total x-ray reflectance fluorescence.

**Results:** In the hippocampus, we observed AUD-related decreased iron, but nothing more. In the substantia nigra, we observed a significant increase in iron and a trend toward an in increase in copper content in females and decreased iron and zinc in males. In the caudate we saw a near doubling of iron in females and a significant increase in copper in females and males.

**Conclusions:** These results provide direct evidence that at least part of AUD-related pathophysiology involves the dysregulation of trace metals in the brain. These changes are more severe in females, which may relate to extended levels of exposure due to slower alcohol metabolism.

## INTRODUCTION

Among the trace metals that have biological functions in eukaryotes, iron is probably the most important among transition metals. It plays essential roles in cellular respiration in enzyme activity, DNA synthesis, hemoglobin function and others. In the brain, iron is essential for neurotransmitter production and function, and myelin production. Consequently, iron is essential for cognition, affect, and motor activity. Iron concentration in the brain is critical. Several proteins control its accumulation, removal, and action. Too little iron in the brain during development can produce cognitive deficits that are permanent (Lozoff et al., 2013) and in adulthood restless legs syndrome (Earley et al., 2014). Alternatively, brain iron concentrations that are greater than homeostatic concentrations can cause damage by the production of free radicals and oxidative stress. Indeed, nearly all neurodegenerative diseases, including Parkinson’s disease (Kulaszyńska et al., 2024; Ahern et al., 2025), have one thing in common, i.e., iron dysregulation with high iron concentrations. We and others have shown that toxin and toxicant exposure can dysregulate brain iron with increased concentrations (Torres-Rojas et al., 2020; Xiu et al., 2025). Recently, we showed that alcohol consumption in mice can dysregulate brain iron and reduce the mass of the hippocampus (Jones et al., 2021). The authors of a recent article proposed that heavy alcohol consumption in humans could dysregulate iron, leading to dementia (Listabarth et al, 2020).

Considering the hypotheses about alcohol and iron, we obtained brain samples from the New South Wales Brain Tissue Resource Centre (Sutherland et al., 2016). The donors of these brains included those diagnosed with AUD and those without alcohol-related problems. We were able to analyze the concentrations of iron, copper, and zinc.

## MATERIALS AND METHODS

### Subjects

Tissue samples, including, hippocampus, substantia nigra, and caudate, were obtained from frozen brains. About 25 mg of each tissue was collected and shipped to the Memphis laboratory. Table 1 presents the numbers and ages of the donors.

**Table 1.** Number of samples and mean age of donors.

| **Donor** | **AUD female** | **Control female** | **AUD male** | **Control male** |
| --- | --- | --- | --- | --- |
| n | 10 | 10 | 10 | 10 |
| Mean Age | 57 | 56.9 | 60.8 | 61.4 |
| s.d. | 14.24 | 15.27 | 14.72 | 14.91 |

### Tissue preparation and analysis

The tissues were wet-ashed in 400 µl ultra-pure nitric acid and digested 48h at room temperature. The samples were diluted by adding 400 µl distilled water. with gallium (1µg ml^-1^ final concentration) added as the internal standard. Ten µl of the sample was placed on a quartz sample carrier and dried on a hotplate. Technical duplicates were read on a Bruker Instruments (Billerica, Mass.) S2 PicoFox Total Reflection X-ray Fluorescence spectrometer with sub ppb sensitivity. Each sample was read for 500 seconds.

### Data analysis

The PicoFox data were expressed as µg g^-1^ for all metals in the three tissues. The data were analyzed by analysis of variance for a two between-subjects variables (diagnosis, sex) and age as covariate (which was non-significant in all analyses). Because this was a one degree of freedom experiment, no post hoc tests were needed.

## RESULTS

### Hippocampus

Table 2 presents the results for Fe, Cu, and Zn in the hippocampus. The only significant main effect observed was a decrease in Fe in both sexes diagnosed with AUD compared to control (F_1,39_=5.10, p<.04).

**Table 2.** Metals concentration in hippocampus in males and females diagnosed with AUD vs. control.

|  | Female AUD | Female Control | Male AUD | Male Control |
| --- | --- | --- | --- | --- |
| Fe |  |  |  |  |
| Mean | 25 | 27.66 | 24.27 | 30.46 |
| s.e.m. | 1.94 | 1.94 | 1.93 | 2.06 |
| Cu |  |  |  |  |
| Mean | 1.96 | 2.19 | 2.53 | 2.48 |
| s.e.m. | 0.26 | 0.26 | 0.26 | 0.27 |
| Zn |  |  |  |  |
| Mean | 12.02 | 10.32 | 11.54 | 13.59 |
| s.e.m. | 1.37 | 1.37 | 1.37 | 1.46 |

### Substantia Nigra

Table 3 and figure 1 show the results of differential diagnosis for the metals. Neither main effect for diagnosis nor sex were significant; however, the diagnosis X sex interaction was significant for iron (F_1,39_=10.62, p<0.004 and zinc (F_1,39_=6.20, p<0.02). The iron concentration for AUD females was significantly greater than controls, whereas there was a significant decrease in iron and zinc for males but not in females.

**Table 3.** Metals concentrations in substantia nigra in males and females diagnosed with AUD vs. control.

|  | Female AUD | Female Control | Male AUD | Male Control |
| --- | --- | --- | --- | --- |
| Fe |  |  |  |  |
| Mean | 131.03 | 83.01 | 104.63 | 128.56 |
| s.e.m. | 11.45 | 10.92 | 10.87 | 10.88 |
| Cu |  |  |  |  |
| Mean | 9.73 | 9.79 | 8.18 | 9.47 |
| s.e.m. | 0.88 | 0.84 | 0.83 | 0.83 |
| Zn |  |  |  |  |
| Mean | 12 | 10.56 | 10.21 | 12.21 |
| s.e.m. | 0.72 | 0.68 | 0.68 | 0.68 |

**Table 3.** Metals concentrations in caudate in males and females diagnosed with AUD vs. control.

|  | Female AUD | Female Control | Male AUD | Male Control |
| --- | --- | --- | --- | --- |
| Fe |  |  |  |  |
| Mean | 100.64 | 53.4 | 69.48 | 64.04 |
| s.e.m. | 7.473 | 8.34 | 7.74 | 7.80 |
| Cu |  |  |  |  |
| Mean | 4.76 | 3.28 | 3.86 | 2.78 |
| s.e.m. | 0.49 | 0.53 | 0.49 | 0.5 |
| Zn |  |  |  |  |
| Mean | 11.80 | 10.97 | 11.02 | 12.11 |
| s.e.m. | 0.83 | 0.90 | 0.83 | 0.84 |

**Figure 1.**
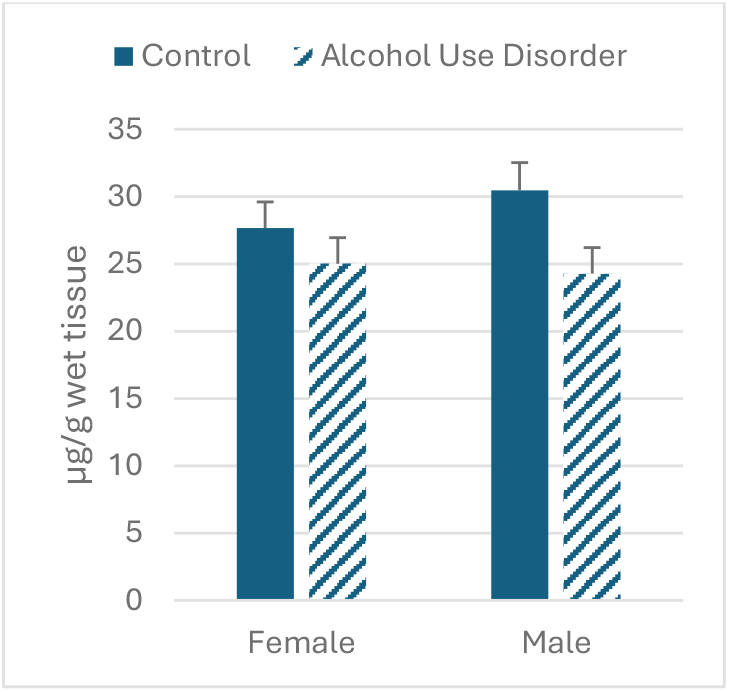
Iron concentration in hippocampus in individuals diagnosed with AUD vs. control.

**Figure 2.**
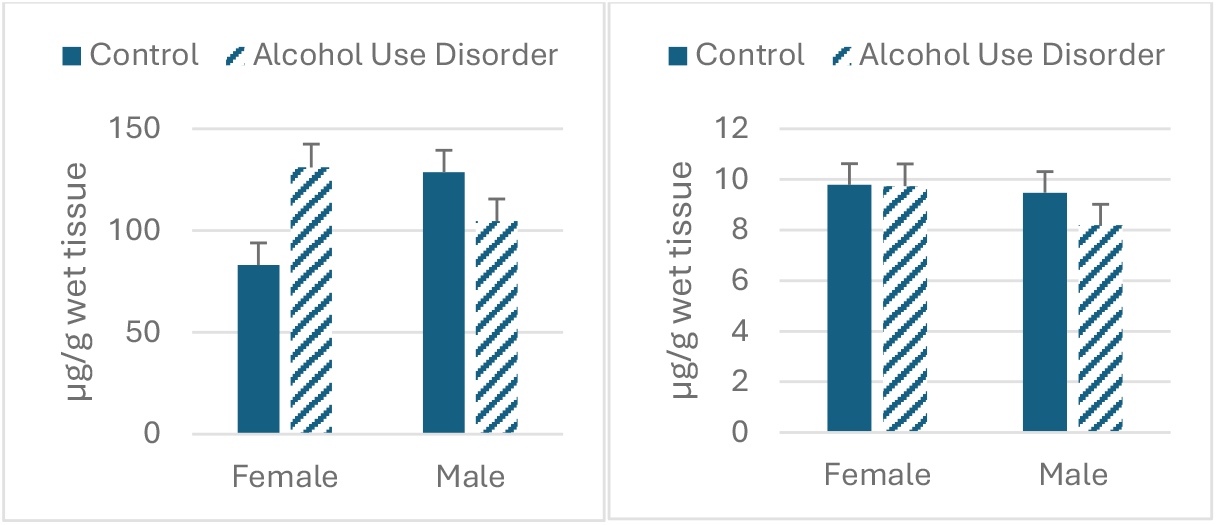
Effect of AUD and sex on substantia nigra iron content (left panel) and zinc (right panel).

**Figure 3.**
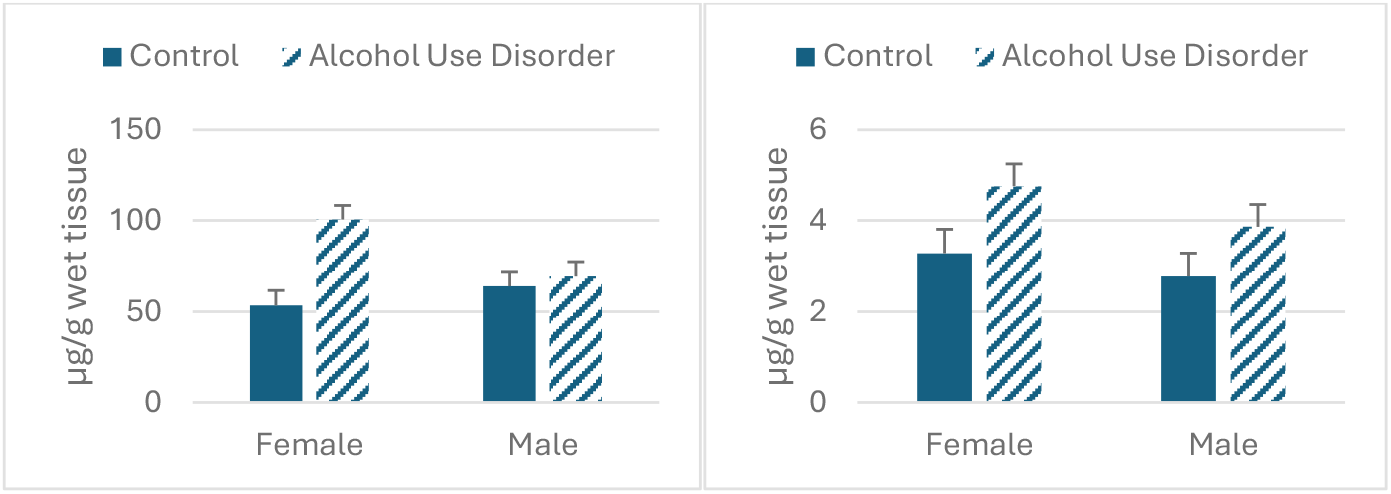
Effect of AUD and sex on caudate iron content (left panel) and copper (right panel).

### Caudate

We observed significant main effects of AUD for iron and copper (F_1,35_=11.24; 6.56, p<0.003,0.02, respectively, We also saw a significant interaction between AUD and sex (F_1,35_=6.96, p<0,02). Iron concentration was increased in both sexes, with a smaller increase in males. Copper was increased in both sexes.

## DISCUSSION

To the best of our knowledge, this is the first study for which the actual concentrations of the metals were measured and reported. The three other studies that reported brain iron susceptibility using magnetic resonance imaging (MRI and quantitative susceptibility mapping), an indirect estimate of amount of iron, in individuals who used alcohol in varying amounts. Juhás and colleagues (2017) reported AUD-related increased susceptibility in caudate, putamen/globus pallidus, and dentate nucleus. In 2022,Topiwala and colleagues measured brain iron in individuals participating in the UK Biobank. They were able to assess the relationship between brain iron content in individuals who reported the amount of alcohol that they normally consumed and not necessarily diagnosed with AUD. They reported alcohol-related increases in MRI quantitative susceptibility at even moderate consumption in serum ferritin and in liver. In the brain, they reported alcohol-related decreases in susceptibility in the thalamus but increases in the substantia nigra and putamen. Adams and colleagues (2023) reported peripheral and central susceptibility in individuals diagnosed with AUD. They reported higher susceptibility in serum ferritin. Considering whole brain susceptibility, they reported increased concentrations in older individuals and in males; however, neither AUD, serum ferritin nor age was associated with overall increases in susceptibility.

Regionally, they reported increased susceptibility in the left globus pallidus. Remarkably, in none of the three studies was sex shown as important.

While MRI imaging of brain iron has its uses, the measurement approximates the actual content. Nevertheless, there is considerable agreement with what we found with direct measurement of brain iron. Although there are some differences in reported structures, all three studies and ours show the striatum to be susceptible to AUD-related increases in iron. Also, our results agree with the Topiwala study (2022) that includes the substantia nigra as well.

The remaining question is exactly how increased iron causes the neurological damage associated with AUD. One likely mechanism is iron-induced neuroinflammation (Ward et al., 2022; Wilcockson and Roy, 2024). Exposure to toxins and toxicants can activate brain microglia and increase proinflammatory cytokines leading to neurological damage. We have shown that treatment with paraquat (Torres-Rojas et al., 2020) and MPTP (Jones et al., 2013) can dysregulate iron in the midbrain and striatum which raises the question whether the toxicity is caused by the agent or by the increased iron. The same question can be addressed to alcohol.

### What about regional differences?

We showed that the nigrostriatal system was particularly vulnerable to AUD-based iron dysregulation. This is interesting because Goutaudier and colleagues (2022) showed in rats that alcohol consumption reduced dopamine activity in the anterior dorsolateral striatum, dorsomedial striatum, and nucleus accumbens, accompanied by increased alcohol consumption. The concept is that dopamine insufficiency in the nigrostriatal system increases alcohol consumption in a feed-forward manner. Moreover, iron and dopamine can combine to produce a toxic redox compound and this is particularly pronounced in the dopamine-rich nigrostriatal system (Hare & Double, 2016).

### Alcohol-related risks: gender differences

Particularly noteworthy is the fact that women are at greater risk for alcohol-related disorders than men. These include liver, cardiovascular, pancreatic, neurological, psychiatric and more (White, 2020). There are several reasons for this; however, two predominant are differences in volume of distribution and in alcohol dehydrogenase (ADH). Concerning volume of distribution, men and women of equal body mass and consuming equal amounts of alcohol, women will experience a higher blood alcohol content. Alcohol distributes to the water compartment and women have a smaller water compartment than men. Men produce more liver and stomach ADH than women and women produce very little stomach ADH. This means that women metabolize alcohol more slowly than men (Chrostek et al., 2003). Here, we show that women are more susceptible to iron dysregulation and possible neuropathology in the nigrostriatal system than men.

### What about copper and zinc?

While the main target of this research was iron, our technical methods allowed us to measure other elements simultaneously, specifically, copper and zinc. Copper is an essential micronutrient for brain development and function. It engages in mitochondrial activity, catecholamine synthesis, myelination and other functions(An et al., 2022). Like iron, it exists in two oxidation states, Cu^+^ and Cu^2+^, the latter being more active chemically and potentially more toxic when out of homeostatic control. Indeed, copper dysregulation is associated with impaired cognition and neurodegenerative diseases and psychiatric disorders (Feng et al.,2023; Gout et al., 2023). Here, we show an increase in copper in the caudate in individuals diagnosed with AUD compared to controls.

Zinc is also an important trace element. It takes part in neuroimmune actions, is a crucial element in transcription factors and is an antioxidant. Its actions in transcription factors takes part in the activities of numerous enzymes important for neuronal metabolism and regulates synaptic activity. Like iron and copper, zinc requires tight regulation. In the brain, zinc deficiency is associated with dementia and other cognitive difficulties. Overload of zinc is associated with neurodegeneration (Skalny et al., 2017; Granzotto et al., 2020; Li et al, 2022). AUD has been shown to interfere with intestinal zinc absorption (Pavuluri er al., 2022) and presumably to reduce zinc systemically and in the brain. The impact of this decreased zinc absorption on regional distribution of brain zinc, is unknown.

## CONCLUSION

Here, we report the direct assessment of brain iron, copper, and zinc in individuals diagnosed with AUD. Accordingly, we show differences in these metals’ response by tissue and by sex/gender. Our work reinforces the importance of alcohol-related iron dysregulation but now includes copper and zinc as important players. This research should open avenues toward increased research into AUD effects on trace metals and related therapeutics.

## ACKNOWLEDGEMENTS

The authors thank Daming Zhuang for her expertise in the analysis of brain samples.

Tissues were received from the New South Wales Brain Tissue Resource Centre at the University of Sydney which is supported by the University of Sydney. Research reported in this publication was supported by the National Institute of Alcohol Abuse and Alcoholism of the National Institutes of Health under Award Number R28AA012725 The content is solely the responsibility of the authors and does not represent the official views of the National Institutes of Health.

## ETHICS STATEMENT

All tissue donors have been de-identified and the Institutional Review Boards of the University of Sydney and the University of Tennessee Health Science Center approved of this project.

## DATA AVAILABILITY STATEMENT

All data can be made available by contacting the corresponding author

## AUTHOR CONTRIBUTIONS

BCJ conceptualized the study and wrote all drafts

WZ directed the analyses, performed the statistical analyses and produced the graphs

GTS performed major copy editing and suggestions for the discussion

JS supplied the tissues and performed copy editing.

## REFERENCES

Adams AR, Li X, Byanyima JI, Vesslee SA, Nguyen TD, Wang Y, Moon B, Pond T, Kranzler HR, Witschey WR, Shi Z, Wiers CE. (2023) Peripheral and Central Iron Measures in Alcohol Use Disorder and Aging: A Quantitative Susceptibility Mapping Pilot Study. Int J Mol Sci. 24:4461.

Ahern J, Boyle ME, Sugrue L, Andreassen O, Dale A, Thompson WK, Fan CC, Loughnan R. (2025) Dietary Factors Affect Brain Iron Accumulation and Parkinson’s Disease Risk. medRxiv [Preprint] 13.24304253.

An Y, Li S, Huang X, Chen X, Shan H, Zhang M. (2022) The Role of Copper Homeostasis in Brain Disease. Int J Mol Sci. 23:13850.

Chrostek L, Jelski W, Szmitkowski M, Puchalski Z (2003) Gender-related differences in hepatic activity of alcohol dehydrogenase isoenzymes and aldehyde dehydrogenase in humans. J Clin Lab Anal. 17:93–6

Earley CJ, Connor J, Garcia-Borreguero D, Jenner P, Winkelman J, Zee PC, Allen R. (2014) Altered brain iron homeostasis and dopaminergic function in Restless Legs Syndrome (Willis-Ekbom Disease). Sleep Med. 11:1288–301.

Gout D, Zhao Y, Li W, Li X, Wan J, Wang F. (2023) Copper neurotoxicity: Induction of cognitive dysfunction: A review. Medicine (Baltimore). 102:e36375.

Goutaudier R, Joly F, Mallet D, Bartolomucci M, Guicherd D, Carcenac C, Vossier F, Dufourd T, Boulet S, Deransart C, Chovelon B, Carnicella S. (2023) Hypodopaminergic state of the nigrostriatal pathway drives compulsive alcohol use. Mol Psychiatry. 28:463–474.

Granzotto A, Canzoniero LMT, Sensi SL (2020). A Neurotoxic Ménage-à-trois: Glutamate, Calcium, and Zinc in the Excitotoxic Cascade. Front Mol Neurosci. 26:13:600089

Hare DJ and, Double KL. (2016) Iron and dopamine: a toxic couple. Brain. 139(Pt 4):1026–35.

Jones BC, Miller DB, O’Callaghan JP, Lu L, Unger EL, Alam G, Williams RW. (2013) Systems analysis of genetic variation in MPTP neurotoxicity in mice. Neurotoxicology. 37:26–34.

Jones BC, Erikson KM, Mulligan MK, Torres-Rojas C, Zhao W, Zhuang D, Lu L, Williams RW. (2021) Genetic differences in ethanol consumption: effects on iron, copper, and zinc regulation in mouse hippocampus. Biometals. 34:1059–1066.

Juhás M, Sun H, Brown MRG, MacKay MB, Mann KF, Sommer WH, Wilman AH, Dursun SM, Greenshaw AJ. (2017) Deep grey matter iron accumulation in alcohol use disorder. Neuroimage. 148:115–122.

Li Z, Liu Y, Wei R, Yong VW, Xue M. (2022) The Important Role of Zinc in Neurological Diseases. Biomolecules. 13:28.

Listabarth S, König D, Vyssoki B, Hametner S. (2020) Does thiamine protect the brain from iron overload and alcohol-related dementia? Alzheimers Dement. 16:1591–1595.

Lozoff B, Smith JB, Kaciroti N, Clark KM, Guevara S, Jimenez E. (2013) Functional significance of early-life iron deficiency: outcomes at 25 years. J Pediatr. 163:1260–6.

Kulaszyńska M, Kwiatkowski S, Karolina Skonieczna-Żydecka, K. (2024) The Iron Metabolism with a Specific Focus on the Functioning of the Nervous System. Biomedicines. 12:595.

Skalny AV, Skalnaya MG, Grabeklis AR, Skalnaya AA, Tinkov AA. (2018) Zinc deficiency as a mediator of toxic effects of alcohol abuse. Eur J Nutr. 57:2313–2322

Pavuluri P, Jangili S, Ryakam L, Vadakedath S, Tummalacharla SC, Kondu D, Kandi V. (2022) The Activities of Zinc and Magnesium Among Alcohol Dependence Syndrome Patients: A Case-Control Study From a Tertiary Care Teaching Hospital in South India. Cureus. 14:e24502.

Sutherland GT, Sheedy D, Stevens J, McCrossin T, Smith CC, van Roijen M, Kril JJ. (2016) The NSW brain tissue resource centre: Banking for alcohol and major neuropsychiatric disorders research. Alcohol. 52:33–39.

Topiwala A, Wang C, Ebmeier KP, Burgess S, Bell S, Levey DF, Zhou H, McCracken C, Roca-Fernández A, Petersen SE, Raman B, Husain M, Gelernter J, Miller KL, Smith SM, Nichols TE. (2022) Associations between moderate alcohol consumption, brain iron, and cognition in UK Biobank participants: Observational and mendelian randomization analyses. PLoS Med. 19:e1004039.

Torres-Rojas C, Zhuang D, Jimenez-Carrion P, Silva I, O’Callaghan JP, Lu L, Zhao W, Mulligan MK, Williams RW, Jones BC. (2020) Systems Genetics and Systems Biology Analysis of Paraquat Neurotoxicity in BXD Recombinant Inbred Mice. Toxicol Sci. 176:137–146.

Ward RJ, Dexter DT, Crichton RR. (2022) Iron, Neuroinflammation and Neurodegeneration. Int J Mol Sci. 23:7267.

White AM. (2020) Gender Differences in the Epidemiology of Alcohol Use and Related Harms in the United States. Alcohol Res. 40:01.

Wilcockson TDW and Roy S. (2024) Could Alcohol-Related Cognitive Decline Be the Result of Iron-Induced Neuroinflammation? Brain Sci. 14:520.

Xiu M, Liu Y, Wang Z, Zhang J, Shi Y, Xie J, Shi L. (2025) Abnormal iron metabolism in the zona incerta in Parkinson’s disease mice. J Neural Transm (Vienna). 132:845–857.

